# Internal bias controls dopamine perceptual decision-related responses

**DOI:** 10.1101/431387

**Authors:** Stefania Sarno, Manuel Beirán, José Vergara, Román Rossi-Pool, Ranulfo Romo, Néstor Parga

## Abstract

Dopamine neurons produce reward-related signals that regulate learning and guide behavior. Prior expectations about forthcoming stimuli and internal biases can alter perception and choices and thus could influence dopamine signaling. We tested this hypothesis studying dopamine neurons recorded in monkeys trained to discriminate between two tactile frequencies separated by a delay period, a task affected by the contraction bias. The bias greatly controlled the animals’ choices and confidence on their decisions. During decision formation the phasic activity reflected bias-induced modulations and simultaneously coded reward prediction errors. In contrast, the activity during the delay period was not affected by the bias, was not tuned to the value of the stimuli but was temporally modulated, pointing to a role different from that of the phasic activity.

## Introduction

The phasic activity of dopamine (DA) neurons codes for reward prediction errors (RPEs) (Schultz et al., 1997; Bayer and Glimcher, 2005; Steinberg, et al., 2013). This is true even under uncertain stimulation conditions (Sarno et al., 2017; Lak et al., 2017; Starkweather et al., 2017). In decision-making tasks the phasic activity reflects internal choice processes (Nomoto et al., 2010; de Lafuente and Romo, 2011), temporal processing (Sarno et al., 2017) and exhibits beliefs about the state of the environment (Starkweather et al., 2017; Sarno et al., 2017; Lak et al., 2017). Importantly, DA responses also depend on the confidence that the animal has on its choice (de Lafuente and Romo, 2011; Sarno et al., 2017; Lak et al., 2017). A crucial aspect of the tasks employed in these studies is the difficulty of a trial, which is usually controlled by the physical features of the stimulus and determines the animal’s choices and decision confidence. However, internally generated biases can also influence the formation of the decision (Körding, 2007; Preuschhof et al., 2010). When biases are present, the difficulty introduced by the experimentalist through the stimulus set can be quite different from the true difficulty experienced by the animal when making a decision. A consequence of this is that its choices and confidence could depend on variables responsible for the internal bias. We then hypothesized that if midbrain DA neurons received information about cortical computations of choice and confidence, their firing activity should be modulated by biases and prior expectations. Besides, if DA neurons estimated RPEs, the bias dependence should manifest itself in a way compatible with that function (Figure 1A).

**Fig. 1:**
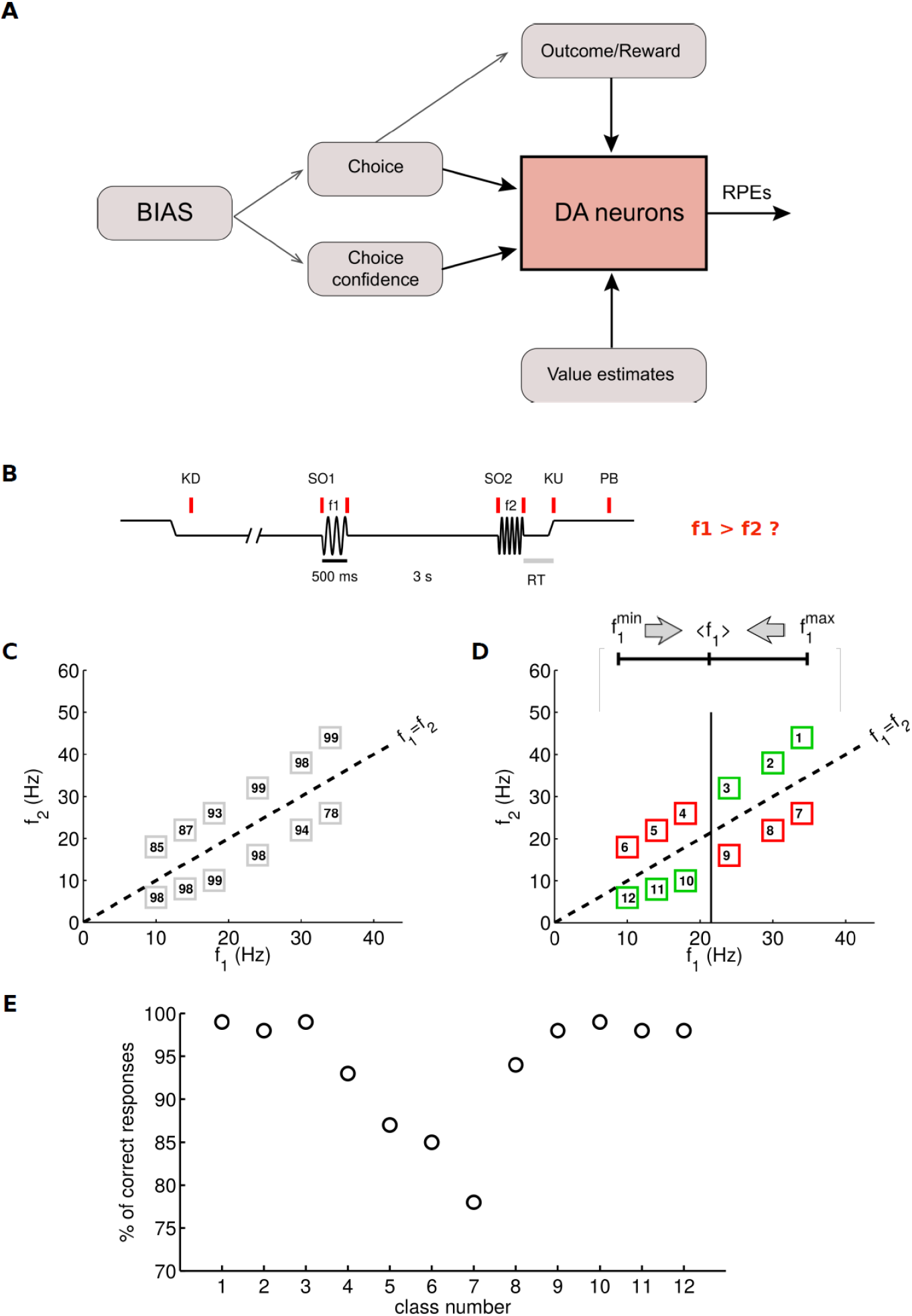
Discrimintaion Task and Contraction Bias. **(A)** Schematic of the influence of the contraction bias on DA neurons and RPEs. The computations of choice and confidence, carried out in cortical circuits, are affected by an internal bias. We hypothesized that if DA neurons receive information about those processes their phasic activity has to reflect the effects of the bias. Besides, if the DA activity represents estimates of RPEs their dependence on the bias has to be consistent with that function. **(B)** Trials began when a mechanical probe was lowered (probe down, PD). The monkey reacted by placing its free hand on an immovable key (key down, KD). After a variable period (1.5–3.0 s) the probe oscillated for 0.5 s at the base (f1) frequency (onset of the first stimulus, SO1). A second vibrotactile stimulus at the comparison frequency (f2) was presented 3 s after the offset of the first stimulus (onset of the second stimulus, SO2). At the end of the second stimulus the monkey released the key (key up, KU) and pressed one of two push-buttons (PB) to indicate whether the comparison frequency was higher or lower than the base frequency. At the PB event, the animal was rewarded for correct responses. **(C)** Stimulus set (i.e., the pairs of frequencies *f*_1_, *f*_2_)) used in the task. Numbers inside the boxes indicate the percentage of correct trials in each class. **(D)** Effect of the contraction bias on the classes in the stimulus set. The upper part of the panel depicts the range of the frequency *f*_1_, from its minimum to its maximum values. As indicated by the two arrows, *f*_1_ is perceived as closer to its mean value in the set, *<f*_1_> (vertical line in the lower part of the panel). Classes for which the bias favors (disfavors) correct responses are indicated by green (red) squares. The numbers inside the squares label the classes following the effect of the contraction bias along each of the two diagonals in the set. **(E)** The accuracy data in panel (C) here are presented versus class number.

An appropriate experimental paradigm to investigate these issues is the 2-interval forced-choice (2IFC) task in which the animal discriminates a physical feature after observing two stimuli presented sequentially, with a delay period between them lasting a few seconds (Green and Swets, 1966). A relevant feature of 2IFC tasks is that the perception of the first stimulus appears shifted towards the center of its range, an effect that introduces a behavioral bias in the comparison between the two stimuli (Hollingworth, 1910). This so-called contraction bias is observed in many instances of delayed comparison tasks in humans (Ashourian and Loewenstein, 2011; Dyjas et al., 2012; Raviv et al., 2012; Fassihi et al., 2014; Akrami et al., 2018), it is also present in the tactile frequency discrimination task in monkeys (Romo et al., 1999), rodents (Fassihi et al., 2014) and in an auditory version of the same task both in rats and humans (Akrami et al., 2018). Here we investigated whether and how this bias affected the DA firing activity employing a somatosensory discrimination task in which monkeys discriminated between the frequencies of two vibrotactile stimuli delivered to one fingertip (Figure 1B), an instance of the 2IFC paradigm in which both stimuli are selected randomly in each trial (Romo et al., 1999).

## RESULTS

### Tactile Frequency Discrimination Task and Neuron Classification

In the task the animal discriminated between the two frequencies, *f*_1_ and *f*_2_, in the flutter range (Figure 1B). Each pair of frequencies adopted during a trial identified a class; the stimulus set, that is, all the classes used during the recordings, is illustrated in Figure 1C together with the animal’s accuracy. The animal obtained reward for correctly identifying the higher frequency (Figure 1B). The electrophysiological results were obtained from midbrain DA neurons responding to reward delivery with a positive phasic activation in correct trials and with a pause in error trials (Bromberg-Martin et al., 2010; Morris et al., 2006). These were 25 out of a total of 136 neurons with task-related activity recorded in VTA (Methods). To further characterize the population of selected neurons, we computed the size of the peak of the reward response in correct trials and estimated its distribution (Methods). This distribution turned out to be bimodal, with a clear separation in subpopulations with low- and high response peaks occurring at 15 Hz (Figure 2A). The subpopulations had roughly the same number of neurons (12 and 13 neurons with low- and high response peak, respectively). We then set out to study these two groups separately. In what follows, we will first investigate the neurons with a low-response peak and we will analyze later the other subpopulation.

**Figure 2.**
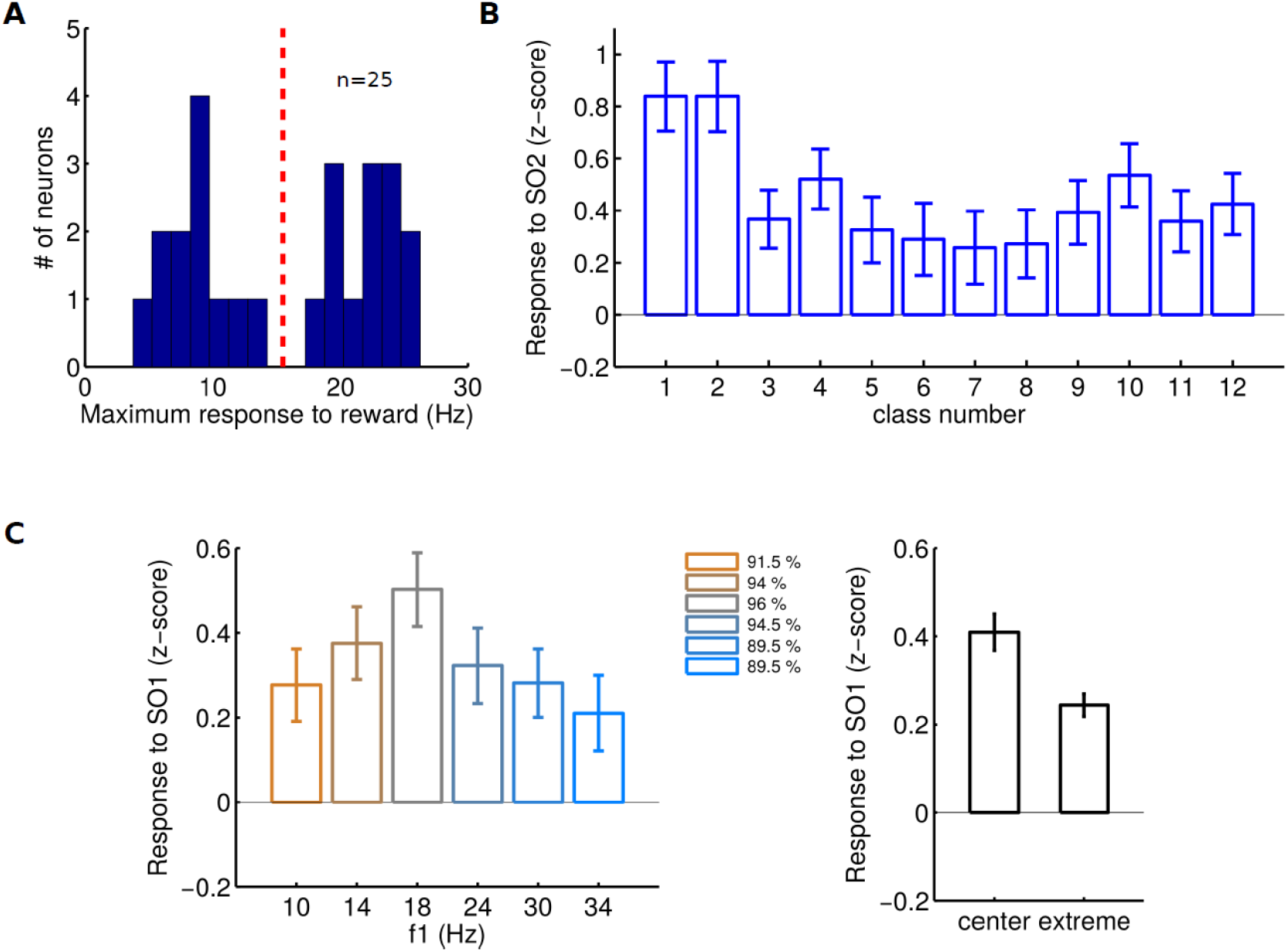
Neuron Classification and Phasic Responses to the Stimuli. **(A)** Fraction of neurons compatible with RL ideas, as a function of their peak response to the reward in correct stimuli. Neurons are organized into two subpopulations with low- and high response peak. The red vertical dashed line indicates the boundary between them at 15Hz. **(B)** Population response (z-score) to the onset of the comparison stimulus (SO2) sorted by stimulus classes. **(C) (Left)** Population response (z-score) to the onset of the base stimulus (SO1) sorted by the frequency of the vibration. Error bars are ± 1 SEM. **Inset:** Percentage of correct responses for each value of *f*_1_. Color code indicates different values of *f*_1_. **(Right)** Population response (z-score) to the onset of the base stimulus for ‘extreme’ values of the base frequency *f*_1_ = 10 Hz and *f*_1_ = 34 Hz) and for ‘center’ values *f*_1_ = 18 Hz and *f*_1_ = 24 Hz). Although the difference in activation did not reach significance value (p = 0.067, two tail t-test), the result of this analysis showed a clear tendency for larger activation when the base stimulus frequency corresponded to trial types more likely to be rewarded. Error bars are ± 1 SEM.

### DA Responses to the Stimuli and Contraction Bias

Performance is only partly controlled by the difference between the two stimulation frequencies, Δ*_f_* = |*f*_1_ − *f*_2_|. In fact, although for most classes Δ*_f_* = 8Hz, the contraction bias induced a dependence of accuracy on the presented class (Figure 1C). Briefly, for classes with correct choice “*f*_1_<*f*_2_“ (upper diagonal) the effect of the bias varies continuously, from favorable to unfavorable, as classes are visited from right to left. For classes with correct choice “*f*_1_>*f*_2_” (lower diagonal) its effect reverts turning more favorable as classes are taken from right to left. We maintained this continuity of the bias effect by labeling classes as in Figure 1D; given this class ordering accuracy is U-shaped, taking the smallest values for classes at the center (Figure 1E).

After the presentation of the second stimulus, the frequency *f*_2_ is compared with the value of *f*_1_ stored in working memory, and cortical neurons readily exhibit the animal’s choice (Hernández et al., 2010). We asked whether the mean DA response to *f*_2_ showed a dependence on the pair (*f*_1_, *f*_2_). For the neuron population with a low response peak, this was indeed the case. When the mean firing rates were displayed vs. class number, the DA phasic responses for classes at the center were less pronounced than for those at the extremes, reflecting the effect of the contraction bias (Figure 2B; one-way ANOVA, p=0.0061).

We next analyzed whether the phasic response to the first stimulus could depend on *f*_1_. Naively, since the trial condition is not fully defined until the application of the comparison stimulus *f*_2_, the DA phasic response to the base frequency is not expected to depend on the value of *f*_1_. Nevertheless, we wondered whether the contraction bias could induce a modulation of the accuracy for fixed values of the first frequency. In the stimulus set, each value of *f*_1_ appears in two classes, one in which the bias favors correct responses and other in which the bias disfavors them (Figure 1C). Hence, it is not *a priori* evident how accuracy at fixed *f*_1_ is affected by the bias. An inspection of the fraction of correct responses for fixed *f*_1_ indicated that the average performance was worse at the end values of *f*_1_ (inset in Figure 2C, left). The DA phasic response to *f*_1_ followed the same trend (Figure 2C, left), exhibiting a slight tendency for a stronger activation for frequencies around the center of its distribution (Figure 2C, right). Given that neurons in several cortical areas encode the value of *f*_1_ parametrically (Romo et al., 1999; Hernández et al., 2010) we next tested if DA responses were consistent with an encoding of *f*_1_. However the population activity did not show any significant dependence on the value of this frequency (one-way ANOVA, p = 0.26), indicating lack of tuning to *f*_1_.

We reasoned that in a similar way that the stimulus physical features can affect the confidence that the animal has about its choices (Kiani and Shadlen, 2009; Kepecs et al., 2008), internally generated biases could have an influence on how confident the animal is (Figure 1A). To investigate this issue we used a Bayesian approach in which observation probabilities are combined with prior information to obtain a posterior probability or belief about the state of the world (Knill and Richards, 1994; Jazayeri and Shadlen, 2015; Ma and Jazayeri, 2014). The prior probabilities of *f*_1_ and *f*_2_ were assumed to be uniform. The observations of *f*_1_ stored in working memory and of the applied *f*_2_ were obtained from Gaussian distributions with standard deviations *σ*_1_ and *σ*_2_, respectively. Choices were made according to the largest of the two posterior probabilities: the belief *b_c_*(H) about the state “*f*_1_ >*f*_2_” (Higher) and the belief *b_c_*(L) about the state “*f*_1_ <*f*_2_” (Lower). The fit of the behavioral data with the Bayesian model yielded *σ*_2_ = 3.2 Hz and *σ*_1_ = 5.50 Hz (Figure 3A. See Methods). Since *σ*_1_ > *σ*_2_ the memory of *f*_1_ deteriorated during the delay period. Interestingly, when this happens a Bayesian model of delayed comparison tasks produces the contraction bias, just because for noisier observations the inferred value of the first stimulus relies more on its prior distribution (Ashourian and Loewenstein, 2011).

**Fig. 3.**
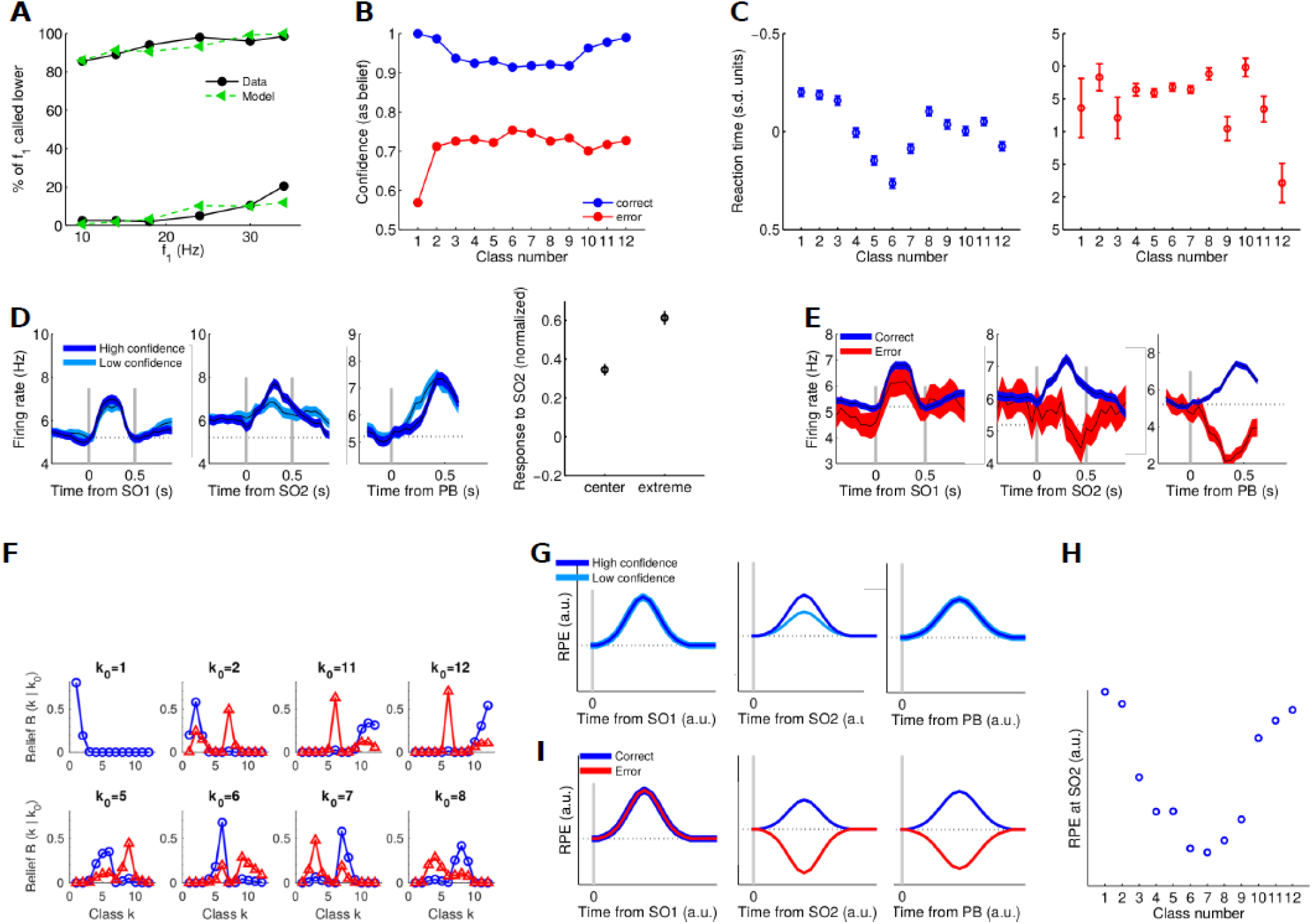
Model-based Analysis of Performance and Confidence and Reinforcement Learning model. **(A)** Model fit of the percentage of times in which the base frequency is called smaller. The behavioral data are indicated with black circles and the model prediction with green triangles. Model best-fit parameter values were *σ*_1_ = 5.34 Hz and *σ*_2_ = 1.75 Hz. **(B)** Confidence, as predicted by the Bayesian model, in correct (blue circles and line) and wrong (red circles and line) trials. **(C)** Normalized response time (RT) in correct (left) and wrong (right) trials as a function of class number (see Methods). **(D)** DA population responses to the two stimuli and to reward delivery in correct trials of two different confidence levels. **Left**: Population firing aligned to the onset of the base frequency (black line, ± SEM colored band) plotted as a function of time for correct trials sorted into two levels of confidence. The gray vertical lines (from left to right) indicate the onset and the offset of the stimulus. **Center**: Same as in the left panel, but with trials aligned to the onset of the comparison stimulus. **Right**: Population firing response aligned to the push button (which coincides with reward delivery), indicated by the gray vertical line (black lines, ± SEM colored bands) plotted as a function of time for correct trials sorted into two confidence levels. **Far right**: Mean response of the DA neurons after the second stimulus for classes with high and low confidence levels. The activation was significantly higher at high-confidence classes (p < 0.05, two tail t-test). **(E)** DA population responses to the stimuli and to reward delivery in correct and error trials. **Left**: Population firing rate aligned to the onset of the base stimulus, for correct and error trials (black line, ± SEM colored band). The vertical gray lines (from left to right) indicate the onset and the offset of the first stimulus. **Center**: As in the left panel, but with trials aligned to the onset of the comparison stimulus. **Right**: As in the left panel but with trials aligned to the push button (reward time), as indicated by the vertical line. **(F)** Distribution of the belief *B*(k|k_0_) that class k has been presented when the true class was k_0_. **Top**: For presented classes favored by the contraction bias (k_0_ = 1, 2, 11, 12). **Bottom**: For presented classes disfavored by the bias (k_0_ = 5, 6, 7, 8). Blue lines: correct trials; red lines: error trials **(G)** Reinforcement Learning model. RPE in correct trials sorted by their confidence level. **Left**: The RPE after the onset of the first stimulus does not depend on the confidence level. **Center**: The RPE after the onset of the second stimulus depends on choice confidence. **Right**: The RPE generated by the model after the reward delivery peaks at a value independent of the confidence level. **(H)** RPE as a function of class number in correct trials. **(I)** RPE as a function of time in correct and error trials. **Left**: The RPE generated by the model after the onset of the first stimulus is similar in correct and error trials. **Center**: The RPE generated by the model after the onset of the second stimulus shows an activation in correct trials and a depression in error trials. **Left**: The RPE generated by the model after the reward delivery.

### Choice Confidence is Affected by the Contraction Bias

Confidence, the probability that for given observations the choice made in a trial is correct, was obtained as the largest of the two beliefs, *b*_c_(L) or *b*_c_(H). We also evaluated the class confidence, the average of confidence over trials of a given class. Class confidence in correct trials vs. class number was U-shaped (Figure 3B, blue line) decreasing (increasing) as the bias was less (more) favorable to making a correct choice. Confidence in the three classes with a different value of Δ*_f_* (classes 1, 11 and 12) are interpreted as follows: class 1 is favored by the bias and by its larger difference Δ*_f_* = 10 Hz, resulting in a large confidence. Classes 11 and 12 are also favored by the bias, but have Δ*_f_* < 8Hz (6Hz and 4Hz, respectively); this increase in task difficulty somewhat compensates the bias-related improvement and confidence in each of these two classes remains smaller than that in class 1. In wrong trials confidence had an inverted-U shape (Figure 3B, red line). The contraction bias modulated confidence in a way similar to how an odor mixture modulated the choice confidence of rodents executing an odor categorization task (Kepecs et al., 2008). But there was a crucial difference: while in the categorization task confidence varied with the physical properties of the stimuli, in the current task it was modulated mainly by an internally generated process. To further support this dependence of confidence on the bias, we reasoned that the response time (RT, defined as the time elapsed from the PU event to the moment when the monkey released his free hand from the key; see Figure 1B) should also reflect the animal’s confidence. If so, in correct (wrong) trials the RT should vary with class number becoming slower (faster) for classes disfavored (favored) by the contraction bias. Figure 3C shows that this was indeed the case (see also Methods).

To analyze how confidence affected the DA activity we divided the set of classes into a high-confidence group, containing classes {1–3, 10–12}, and a low-confidence group, with classes {4–9} (Figure 3B). During the presentation of the base stimulus the firing rate in low-confidence classes was not significantly different than in high-confidence classes (p = 0.58, two tail t-test; Figure 3D, left). During the comparison period the DA firing rate reflected the animal’s confidence level (Figure 3D, center) and the responses of high-confidence classes were significantly higher than those of low-confidence classes (p < 0.05, two tail t-test) (Figure 3D, far right). Approximately 250 ms after the onset of the comparison stimulus the firing rates of the two groups detached from each other, with high-confidence classes having a higher value (p < 0.05; sliding ROC analysis with permutation test. See Methods). This dependence on confidence while the decision is being formed also appeared in other experiments (Sarno et al., 2017; Lak et al., 2017) but, again, the crucial difference is that in the discrimination task this effect was generated by an internal bias. The phasic activity after the delivery of the reward peaked at the same value for both confidence levels (see Figure 3D, right), a result that implies that the activity depended neither on Δ*_f_* nor on the bias.

Phasic responses to the first stimulus in correct and error trials had a similar temporal profile (Figure 3E, left). In contrast, the application of the second stimulus produced quite different responses; after some latency the activity in wrong trials deviated from the activity in trials with correct choices (Figure 3E, center). The number of error trials was too small to investigate the effect of the confidence level on the pause in activity. At reward delivery the DA phasic activity showed a typical temporal pattern, reducing its firing rate in error trials and increasing it transiently in correct ones (Figure 3E, right); in contrast to the responses to the second stimulus, the trends of activation in correct and error trials departed from each other soon after the delivery of the reward.

The above results about confidence (Figure 3B) and the phasic DA responses (Figs. 3D, E) can be interpreted in the framework of Bayesian decision theory. The difference between high- and low-confidence classes arises from the dispersion of the belief that class k has been presented when the true class was k_0_ (denoted as *B*(k|k_0_), see Methods). Specifically, low-confidence correct trials occur because the belief *B*(k|k_0_) is spread over classes k of the wrong choice, due to noisy observations and to the internal bias. Importantly, the bias contributes to this effect because it tends to form beliefs in classes other than the true one. In fact, the spread of *B*(k|k_0_) over classes of the wrong choice becomes more important if, for the presented class k_0_, the effect of the contraction bias is against reporting the correct choice (blue lines in Figure 3F, bottom). Then confidence -the largest of the total beliefs *b*_c_(H) or *b*_c_(L)- is around its minimum (Figure 3B). Under such conditions decisions are difficult and therefore the onset of the second stimulus generates a low prediction of reward (Figure 3D). The opposite is true for classes favored by the bias: the dispersion of the belief is small and is constrained mostly to classes leading to the correct choice (blue lines in Figure 3F, top). For these classes decisions are easier and the onset of the second stimulus produces a higher response. Error trials occur when the total belief about the wrong choice is larger than 0.5. For presented classes disfavored by the contraction bias, the belief *B*(k|k_0_) spreads over several classes of both choices and confidence cannot be large (red lines in Figure 3F, bottom). Even so, for those classes confidence reaches its highest values in wrong trials (Figure 3B, red line). This is because for presented classes favored by the contraction bias errors are mainly due to rare noisy observations and the belief *B*(k|k_0_) remains concentrated in a single wrong class (red lines in Figure 3F, top); confidence about the (wrong) choice is then closer to 0.5.

### A Reinforcement Learning Model Based on Belief States Describes Well the Effect of the Bias on the DA Phasic Responses

To test whether the phasic responses can be attributed to dopamine RPEs we constructed a reinforcement learning model and checked if it was able to reproduce the observed responses. The model consisted in a temporal difference (Sutton and Barto, 1998) module that received beliefs estimated by the same Bayesian model used above (Methods). As in the DA phasic responses (Figure 3D), the RPE after the application of *f*_1_ was independent of the confidence level (Figure 3G, left) and the same happened after the delivery of reward (Figure 3G, right). Instead, after the second stimulus it was modulated by the level of confidence (Figure 3G, center). Furthermore, the RPE estimated for individual classes followed a U-shaped profile (Figure 3H), as the DA responses (Figure 2A). The RPE generated after the three relevant task events in correct and error trials (Figure 3I) was also consistent with the data (Figure 3E).

### The Activity during the Delay Period is Positively Modulated and not Tuned to the Stimulation Frequency

During the delay period neurons in several prefrontal areas exhibit a temporally-modulated persistent activity, tuned parametrically to the value of *f*_1_ (Romo et al., 1999; Brody et al., 2003; Hernández et al., 2010). We addressed several questions about the activity of DA neurons during that period: Is it affected by the contraction bias? Is it temporally modulated during that period? Do neurons exhibit tuning to the value of *f*_1_? Do they code reward-related information? Figure 4A shows the temporal profile of the trial-averaged spiking activity until the end of the delay period for two example cells. The firing rate of the first neuron was modulated in time, increasing throughout the duration of the delay period (Figure 4A, left). Instead, the activity of the other neuron remained closer to its baseline value (Figure 4A, right). At the population level the activity showed a positive modulation throughout the whole delay period (Figure 4B, left), a behavior that is not compatible with coding of RPEs. The mean population firing rate started to increase immediately after the offset of the first stimulus and did so until the application of the second stimulus (Figure 4B, right). A previous work that investigated the DA responses in monkeys executing a visual search task that required working memory did not observe neurons with an ascending profile (Matsumoto and Takada, 2013). The different conclusion could be due to differences in the cognitive requirements of the two tasks or in the recordings sites.

**Figure 4:**
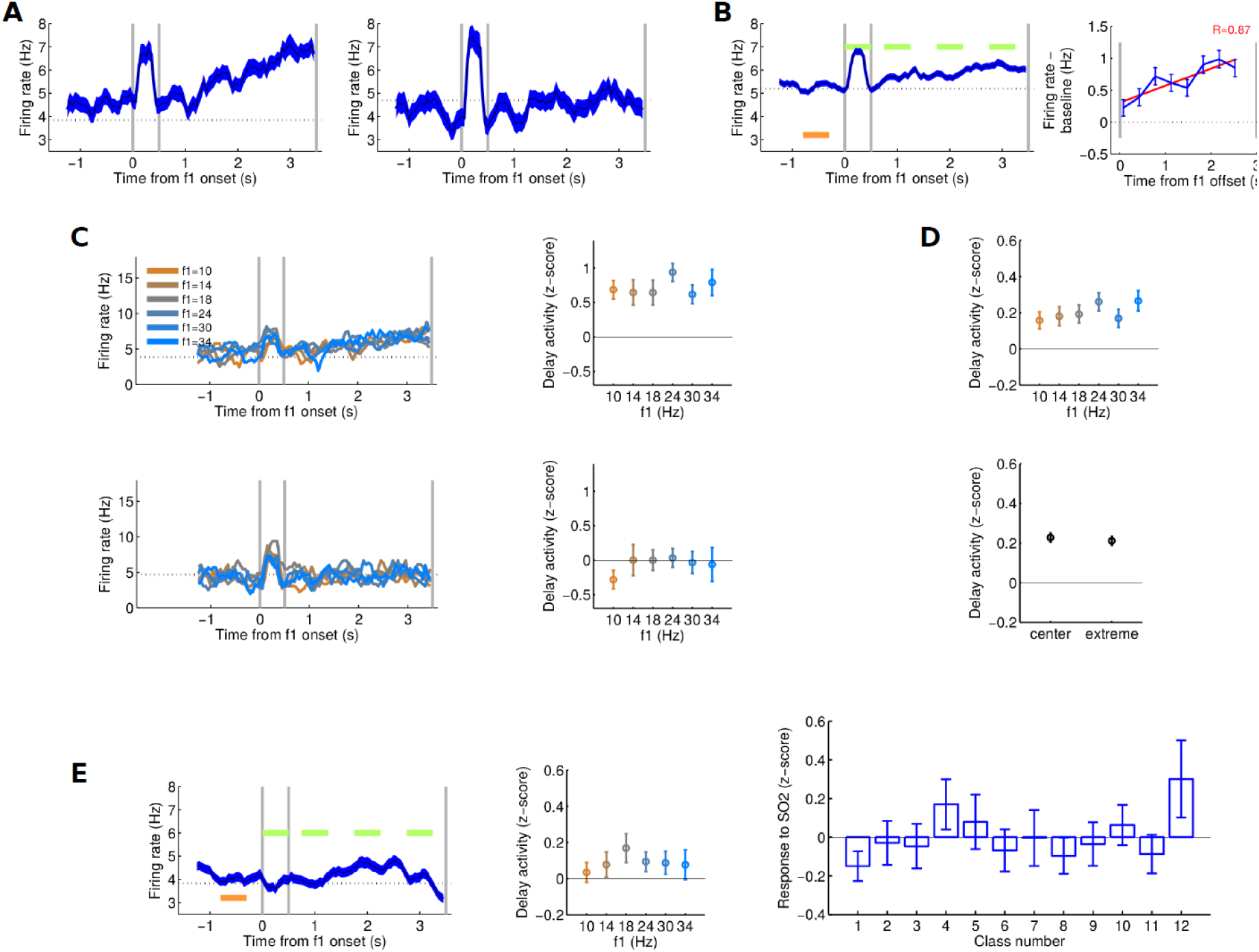
DA Activity during the Delay Period. **(A)** Firing rate for two example neurons plotted as a function of time for rewarded trials. Trials are aligned to the onset of the base stimulus (black line, ± 1 SEM colored band). The gray bars, from left to right, indicate the onset and the offset of the first stimulus, and the onset of the second stimulus. **(B) Left**: Population firing rate plotted as a function of time for rewarded trials (black line, ± 1 SEM colored band). Trials are aligned to the onset of the base stimulus. The gray bars (from left to right) indicate the onset and the offset of the first stimulus and the onset of the second stimulus. Two analyze whether the population firing rate was affected by the first stimulus and how it was modulated during the delay period, we compared the mean firing rates estimated in four portions of its timecouse (green horizontal bars) with the mean firing rate during a time segment previous to the application of the first frequency (red bar). The mean firing of the four temporal segments showed significant positive modulations (one tailed Wilcoxon signed-rank test; p=0.001, p= 0.001, p< 0.001 and p<0.001, respectively) **Right**: Mean population activity of DA neurons during the delay period. Gray vertical lines mark the offset of the base stimulus and the onset of the comparison stimulus. The blue line and symbols are mean-baseline-subtracted firing rate in non-overlapping bins of 350 ms. Error bars are ± 1 SEM. **(C)** Absence of *f*_1_ tuning during the delay period (same two example neurons as in A) **Left**: Firing rate of each of the two neurons sorted according to the value of *f*_1_ in correct trials. Trials are aligned to the onset of the base stimulus. The gray bars (from left to right) indicate the onset and the offset of the first stimulus, and the onset of the second stimulus. **Right**: Mean response (z-score) during the delay period, for the two neurons shown at the left panel, for ‘extreme’ frequencies (*f*_1_ = 10 Hz and *f*_1_ = 34 Hz) and for ‘center’ frequencies (*f*_1_ = 18 Hz and *f*_1_ = 24 Hz). Error bars are ± 1 SEM. **(D)** Absence of *f*_1_ tuning during the delay period **Top**: Mean normalized activity (z-score) of the population sorted according to the value of the first frequency (Methods). Error bars are ± 1 SEM. **Bottom**: Mean population response (z-score) during the delay period for ‘extreme’ frequencies (*f*_1_ = 10 Hz and *f*_1_ = 34 Hz) and for ‘center’ frequencies f_1_ = 18 Hz and *f*_1_ = 24 Hz). Error bars are ± 1 SEM. **(E)** Analysis of the subpopulation with high peak response to the delivery of reward (as defined in Figure 2A). **Left**: Same as in panel B (Left). Two analyze whether the population firing rate was affected by the first stimulus and how it was modulated during the delay period, we compared the mean firing rates estimated in four portions of its timecouse (green horizontal bars) with the mean firing rate during a time segment previous to the application of the first frequency (red bar). There was no significant modulations in the two first segments but the center portion of the delay period exhibited a significant positive modulation and the last segment a negative modulation (one tailed Wilcoxon signed-rank test; p=0.96, p= 0.87, p< 0.05 and p<0.05, respectively) **Center**: Same as in panel D (Top) **Right**: Population response (z-score) to the onset of the comparison stimulus (SO2) sorted by stimulus classes.

We next asked whether this modulation could simply be a reflection of a tuned time-varying signal received from prefrontal inputs. If so, one would expect the firing rate of DA neurons to be tuned to *f*_1_, at least during a portion of that period. We then obtained the temporal profile of the neurons’ firing rates sorting trials according to the value of *f*_1_. For the same two example neurons, the curves for different values of *f*_1_ appeared superimposed (Figure 4C, left) and the temporal averages of the firing activity (z-score) over the whole delay period at fixed *f*_1_ did not exhibit significant differences (Figure 4C; right; one-way ANOVA, p = 0.69 and p = 0.81, for neurons 1 and 2, respectively). This absence of tuning was also observed at the population level during the entire delay period (Figure 4D, top; one-way ANOVA, p = 0.63). Even during the last 500 ms of the delay period, where the activity of neurons in prefrontal areas suggests a recovery of the information about the base frequency (Romo et al., 1999; Hernández et al., 2010; Brody et al., 2003; Rossi-Pool et al., 2016), DA neurons did not exhibit tuning to *f*_1_ (one-way ANOVA, p = 0.71). We next tested the existence of a bias effect; unlike the DA activation after the onset of the base stimulus (Figure 2B, bottom), during the delay period the DA activity did not show any difference between the extreme and the central values of the *f*_1_ distribution (p = 0.95, two tail t-test; Figure 4D, bottom).

### Population with a high peak response to reward delivery

All the previous analyses referred to the subpopulation with a peak response to the reward lower than 15 Hz. The other subpopulation (peak response to the reward higher than 15 Hz) behaved very differently under the application of the two frequencies. It did not exhibit a phasic response to the base frequency (Figure 4E, left) and the (weak) responses to the comparison stimulus were not organized as a U-shaped pattern as a function of class number (Figure 4E, right). Thus, this subpopulation was not affected by the contraction bias and did not code RPEs. On the other hand, its firing activity during the delay period behaved more similarly to the other subpopulation: it was modulated in time (positively modulated during the center portion and negatively modulated towards the end of that period; Figure 4E, left) and it was not tuned to the frequency of the base stimulus (one-way ANOVA, p = 0.83, Figure 4E, center).

## DISCUSSION

Our work provides the first results about the dopamine activity in perceptual decision making when choices are affected by an internal bias. Natural signals tend to vary slowly with time and the brain has adapted to this temporal structure developing circuits able to anticipate future stimuli, e.g., by keeping an average of the recent past inputs (Dyjas et al., 2012; Raviv et al., 2012). In contrast, in the experiment the applied frequencies were selected independently, producing abrupt stimulation changes in consecutive trials. The contraction bias arises from the contradiction between the expected and the received stimuli. Our results suggest that DA neurons processed reward-related information on the basis of signals received from those adapted circuits. Following a model-based approach in which DA neurons estimated RPEs using belief states (Rao, 2010; Sarno et al., 2017; Lak et al., 2017; Starkweather et al., 2017), we found that choices and confidence were greatly controlled by the contraction bias and that DA phasic responses exhibited this modulation during the comparison period. In contrast, the phasic response to the first stimulus and the response to the reward were not affected by confidence.

In fact, the delivery of reward generated a firing response of similar amplitude for both low- and high-confidence classes. This pattern of response is similar to that we encountered previously in a tactile detection task (Sarno et al., 2017). In that case, the response to the reward delivery in hit trials did not depend on the stimulus amplitude, which was the physical parameter that defined the difficulty of the task. However, another recent study that analyzed the response of DA neurons in the random dots motion task showed that the response to a feedback tone, announcing the trial outcome, did depend on the motion coherence (Lak et al., 2017). We speculate that the different pattern of responses could be related to the fact that in the motion discrimination task the stimulus was still present when the animal communicated its decision and received the feedback tone. On the contrary, in the somatosensory detection and discrimination tasks, the relevant stimulus had already disappeared when the animal was rewarded. Therefore, it is possible that in the motion discrimination task the neural activity maintained some dependence on the confidence that determined the decision, whilst in the detection and frequency discrimination tasks this dependence was lost after the offset of the stimulus.

The responses to the second stimulus depended on the confidence level. However, this dependence became significant only after a long latency (Figure 3D, center). Furthermore, the responses in correct and error trials were quite different. While in correct trials the firing activity started to increase shortly after the stimulus onset, in wrong trials it decreased significantly only after a longer latency (Figure 3E, center). These long latencies can be compared with those of cortical signals related to decision processes. Although in some sensory areas (e.g., S1) there are not significant differences between the responses in correct versus error trials, a population of neurons in S2 is tuned to (f1-f2) and distinguishes well between the two trial types after about 220 ms (Romo et al., 2002). Neural populations with activity tuned to the frequency difference and correlated with the animal’s behavior abound in several prefrontal areas (Hernandez et al., 2010). It is plausible that cortical neurons send this information to the DA midbrain system; in turn, these neurons could use those signals to comply with their own function of computing and representing the error on the prediction of the future reward. To quantify the latency of the response to the comparison stimulus in wrong trials, we computed the area under the ROC curve (AUROC) in sliding time windows (see Methods). DA neurons showed a latency of about 260 ms, a value which is slightly longer than the latency of S2 and PFC neurons tuned to (f1-f2) (Hernández et al., 2010). This supports our hypothesis that DA neurons receive signals from cortical areas containing information about choice and confidence (Figure 1A). Previous investigation on a tactile detection task had reached a similar conclusion (de Lafuente and Romo, 2012).

The DA signal during the delay period was quite different from the phasic response to the first stimulus in that it was not affected by the contraction bias. It also differed from the delay activity in prefrontal cortex, where neurons encode parametrically the memorized value of the frequency (Romo et al., 1999). Its only distinctive feature was the existence of neurons with an ascending temporal profile. A signal with these characteristics could fit a suggested role of the DA activity in stabilizing short-term memory in prefrontal cortex (Seamans and Yang, 2004; Kodama and Watanabe, 2017).

Overall our results underline the essential role of internally generated biases in understanding the functional role of DA activity in perceptual decision making tasks.

## METHODS

### Discrimination Task

Monkeys were trained to perform the vibrotactile task as depicted in Figure 1B. Trials began with the probe indenting the finger (Probe Down, PD), followed by the monkey grasping a metal bar with his other hand (Key Down,KD) to signal readiness. After a variable delay of 1500–3000 ms, the first stimulus was applied for 500 ms, followed by a 3000 ms delay and the second stimulus. The monkey then released the bar (Key Up, KU), indicated the choice by pressing one of two push-buttons with the right hand (Push Button, PB), and was rewarded for correctly discriminating the higher frequency. Stimuli were delivered to the skin of the distal segment of one digit of the restrained hand, via a computer-controlled stimulator (BME Systems; 2 mm round tip). Initial probe indentation was 500 μm. Vibrotactile stimuli were mechanical sinusoids. Stimulation amplitudes were adjusted to produce equal subjective intensities.

Animals were handled in accordance with standards of the National Institutes of Health and Society for Neuroscience. All protocols were approved by the Institutional Animal Care and Use Committee of the Instituto de Fisiología Celular (UNAM).

### Recordings

Recordings were obtained with quartz-coated platinum-tungsten microelectrodes (2 to 3 MΩ; Thomas Recording) inserted through a recording chamber located over the central sulcus, parallel to the midline. Midbrain DA neurons were identified on the basis of their characteristic regular and low tonic firing rates (1–10 spikes per second) and by their long extracellular spike potential (2.4 ms ± 0.4 SD). The group of 15 cells used for the analysis corresponded to those neurons that showed a phasic increase in discharge caused by the delivery of reward.

### Analysis of the firing activity

To estimate the temporal profile of the firing rate, for each neuron we counted the number of spikes produced in 300 ms sliding windows shifted every 50 ms.

The responses to the first stimulus (Figure 2C, left) were measured in a 450 ms window centered 280 ms after the stimulus onset. The responses to the second stimulus (Figure 2B) were measured in a 450 ms window centered 320 ms after the stimulus onset. The responses during the delay period (Figures 4C right, 4E center and 4D were measured during its entire duration (3 s). The z-scores of these mean responses were standardized with respect to a temporal window preceding the onset of the base stimulus (the window lasted 500 ms and was centered 1000 ms after the KD).

### Latency analisys

The timecourse of the firing rates of the responses to the second stimulus depend on the confidence level of the trial (Figure 3D) and on whether the trial is correct or not (Figure 3E). To determine the time when two timecourses get apart we applied a receiver operating curve (ROC) analyses in sliding windows. This analysis was done in the period lasting from 300 ms before the onset of the comparison stimulus to 200 ms after its offset. For each neuron we obtained the normalized firing rate (z-score) in sliding windows of 250 ms shifted in 1 ms steps. We used the z-scores of all neurons and trials to calculate the ROC curve at each time bin. The area under the constructed ROC curve (AUROC) was used as the index indicating differential neuronal activity across different trial types. Values of the AUROC higher or lower than 0.5 indicated that, at the population level, one type of trials evoked a higher or lower DA response than the other. To determine the statistical significance of the computed AUROCs, we used a permutation test with 1000 resamples. Significance was assessed when the permutation test indicated statistical significance (p < 0.05) in 50 consecutive time steps. The latency was defined as the time when these 50 consecutive time steps first appear.

### Neuron selection

From the total number of recorded neurons (196) we selected those cells with responses to reward delivery in correct trials significantly higher than the responses to reward omission in wrong trials (P < 0.05, two-sample t test). For each neuron we obtained the maximal response to reward delivery in correct trials. The time of maximal response was assessed by computing the firing rate as a function of time, using 100 ms sliding windows displaced every 100 ms. The activity after reward delivery or omission in correct and error trials was calculated in a window of, respectively, 200 ms and 500 ms, centered at the time of maximum response. The number of neurons compatible with this criterion was n=25.

To further evaluate the selection criterion, we took the 25 selected neurons and shuffled the labels among correct and incorrect trials of each neuron (see, e.g., Romo et al., 1999). We used a permutation test with 1000 resamples to re-analyze the response of these neurons to reward delivery. We found that none of the shuffled neurons resulted reward-responding. Thus, the net probability of marking a neuron as reward-responding by chance was P < 1/25 = 0.04

### Analysis of behavioral data

Behavioral data were obtained on average from 2226 trials per stimulus class. On each session, the response times for each of the two decision outcomes, were standardized (z-score) with respect to the mean and STD (Figure 3C).

### Bayesian model for the discrimination task

The discrimination task was modeled in a Bayesian framework. Prior probabilities of *f*_1_ and *f*_2_ were taken uniform. It was assumed that the animal knew the class structure used in the experiment (Figure 1C) but it had access only to noisy representations (observations) of the two frequencies presented in the trial (denoted by *f*_1,0_ and *f*_2,0_). At the onset of the second stimulus, an observation *o_2_* of the frequency *f*_2,0_ was obtained from a Gaussian distribution with standard deviation *σ*_2_. The observation of the first frequency, taken also at the presentation of the second frequency, had to be taken from the content of working memory and is denoted by *o*_1_^*^. The distribution of this observation was also taken as Gaussian, but with a different standard deviation, *σ*_1_. The belief state B(*c*_k_ | o_1_^*^, o_2_) about the class c_k_ (where k=1, …, 12 labels the classes) was defined as the set of the posterior probabilities 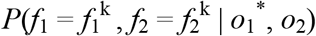 that the class 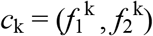 had been presented in the trial, conditioned to the observations *o*_2_ and *o*_1_^*^.

The belief state B(*c*_k_ | *o*_1_^*^, *o*_2_) can be written as:

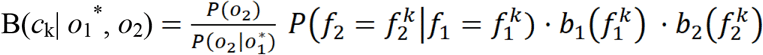

The second factor is a transition matrix relating the first and the second stimulation frequencies. Since we assumed that the animal had perfect knowledge of the class structure, the only non-zero matrix elements correspond to the 12 classes of the experiment. Furthermore, since for a given value of the first frequency the second frequency can only take two possible values, with equal probabilities, all non-zero transition probabilities equal 0.5. The last two factors are the beliefs about the first and second frequencies being those of class k, defined as 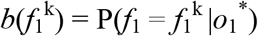 and 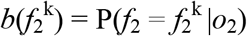.

The sums of the B(k| *o*_1_^*^, *o*_2_)’s over classes k with a fixed choice, give the beliefs about which of the two frequencies is the highest. These two sums were denoted by *b_c_*(H) (belief about the choice “*f*_1_ > *f*_2_”) and *b*_c_(L) = 1 − *b*_c_(H) (belief about “*f*_1_ < *f*_2_”). Choices were made according to the larger of these two beliefs. The performance is measured by the fraction of trials of a given class in which the decision is correct.

The two unknown model parameters, the standard deviations *σ*_1_ and *σ*_2_, were adjusted minimizing the mean squared error between the model performance and the animal’s performance. For the recorded performance data in Figure 1C, the cost function was convex. Using simulated annealing algorithms we found the optimal parameter values, *σ*_1_ = 5.5 Hz and *σ*_2_ = 3.2 Hz.

The confidence *c* of a given trial is *c* = max[*b*_C_(H), 1-*b*_C_(H)], which is bounded between 0.5 and 1. The confidence of a given class was defined as the average of *c* over trials where that class was presented. The confidence in hits (errors) of a given class was defined as the average over the correct (wrong) trials where the class was presented.

Finally, the *class belief state* B(k|k_0_) was defined as the average of the belief state B(k| *o*_1_^*^, *o*_2_) over all trials with a fixed presented class c_k0_ = (*f*_1,0_, *f*_2,0_).

### Reinforcement learning mode

The RL model consists in a Bayesian module, similar to the Bayesian model described above, and a temporal difference (TD) module. The latter consists in an Actor/Critic architecture.

Given the continuous nature of the belief states, value estimation and action selection are implemented using function approximation techniques. Decision rely on a softmax policy that takes the belief state B(ck | o1*, o2) about the class ck and the belief about the choice bc as input. Learning occurs via a TD(λ) algorithm (Sutton and Barto, 1998).

## EXPERIMENTAL PROCEDURES

All protocols were approved by the Institutional Animal Care and Use Committee of the Instituto de Fisiología Celular, Universidad Nacional Autónoma de México.

## ACKNOWLEDGMENTS

This work was supported by Grants FIS2015-67876-P from the Spanish Ministry of Science, Innovation and Universities (to S.S., M.B. and N.P.) and by the Dirección de Asuntos del Personal Académico de la Universidad Nacional Autónoma de México, Consejo Nacional de Ciencia y Tecnología (R.R.) and Fondo Jaime Torres Bodet de la Secretaría de Educación Pública, México (R.R.).

## AUTHOR CONTRIBUTIONS

R.R., J.V. and R.R.-P. performed experiments. S.S., M.B. and N.P. analyzed data and designed the models. S.S. and M.B. wrote the codes. N.P. and S.S. wrote the paper helped by the other authors.

